# Spinosaurids as ‘subaqueous foragers’ undermined by selective sampling and problematic statistical inference

**DOI:** 10.1101/2022.04.13.487781

**Authors:** Nathan Myhrvold, Paul C. Sereno, Stephanie L. Baumgart, Kiersten K. Formoso, Daniel Vidal, Frank E. Fish, Donald M. Henderson

**Author notes:** **Correspondence and requests for materials** should be addressed to Nathan Myhrvold.

## Abstract

Fabbri et al.^1^ claim that the huge sail-backed dinosaur *Spinosaurus aegyptiacus* and the spinosaurid *Baryonyx* were “subaqueous foragers,” diving underwater in pursuit of prey, based on their measure of bone “compactness.” Using thin-sections and computed tomographic (CT) scans of thigh bone (femur) and trunk rib from various living and extinct vertebrates, they claim to be able to distinguish taxa with “aquatic habits” from others. Their conclusions are undermined by selective bone sampling, inaccuracies concerning spinosaurid bone structure, faulty statistical inferences, and novel redefinition of the term “aquatic.”

## Selective bone sampling

A second femur of *Spinosaurus*^2^ (Fig. 1a, b), which is nearly identical in size to the infilled neotypic femur^3^ in their study (Fig. 1c), has a significant medullary cavity lined with cancellous bone that would register as significantly less dense as a thin section at mid shaft. Medullary cavities are also variably present in forelimb bones of *Spinosaurus* (Fig. 1d) resembling those in the long bones of *Suchomimus*, a fully “terrestrial” spinosaurid by their account. Fabbri et al.^1:ED, Fig. 10^ state that *Spinosaurus* and *Baryonyx* “possess dense, compact bone throughout the postcranial skeleton,” yet all three have pneumatic spaces in their cervical column^4^ that exceed in volume the variable long bone infilling, as well as large medullary cavities hollowing the centra at the base of the tail. Neither of these features are present in any secondarily aquatic vertebrate divers that employ bone density as ballast.

**Fig. 1.**
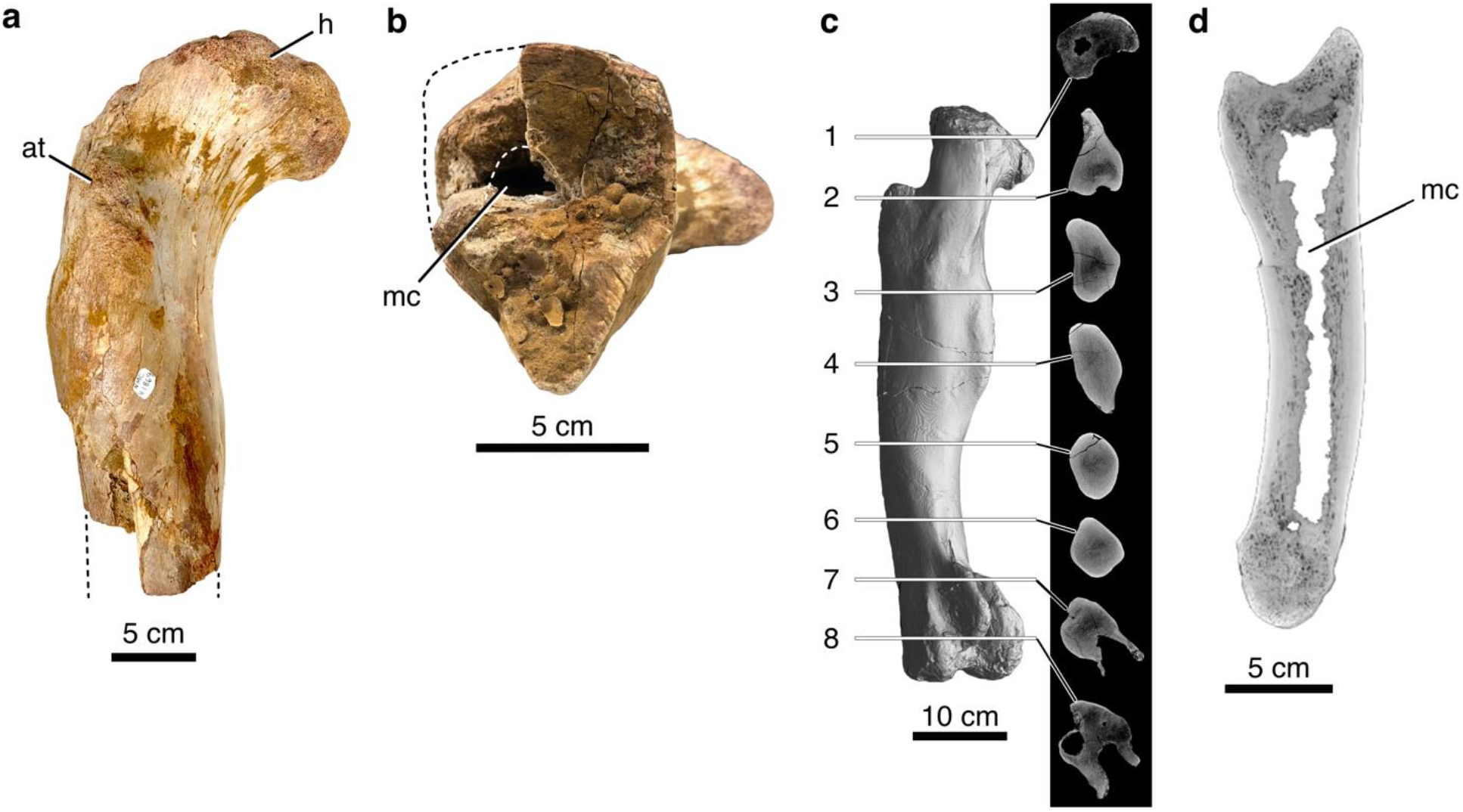
*Spinosaurus cf. aegyptiacus* femur. **a**, Proximal half of the right femur (NMC 41869). **b**, mid-shaft cross-section in distal view. **c**, CT scan of the left femur of the neotype (FSAC-KK 11888) with eight cross sections. **d**, CT scan of right phalanx I-1 in sagittal cross section (UCRC PV8). FSAC, Faculté des Sciences Aïn Chock, University of Casablanca, Morocco; NMC, National Museum of Canada, Ottawa. UCRC, University of Chicago Research Collection, Chicago. at, anterior trochanter, h, head; mc, medullary cavity.

The increase in mass from non-pachystotic bone infill in *Spinosaurus* is less than the mass lost from downsizing its hind limb compared to *Suchomimus* and negligible in a predator weighing at least five metric tonnes. The infilled *Spinosaurus* femur still retains the common bone density gradient (increasing toward minimum shaft diameter; Fig. 1c) for increased resistance to torsion and bending, the usual explanation for non-pachystotic long bone infilling in large facultative bipeds (hadrosaurids) and quadrupeds (elephants, sauropods)^5,6^.

Fabbri et al. use “global bone compactness” (*Cg*) over alternative bone density metrics that have proven better correlated with lifestyle^7^, and they do not detail how they derived that metric from a potpourri of CT scans and physical thin sections. The thin sections for all three spinosaurids in the study come from the distal half of the femur (Fig. 1c, between sections 5 and 7). Fabbri et al.^1:Fig. 1b^ depict the femoral cross section for *Suchomimus* with a larger medullary cavity than is preserved on the original thin section (see Supplementary information), favoring their interpretation of *Suchomimus* as fully “terrestrial” versus *Baryonyx* as “aquatic,” two spinosaurids so similar in skeletal morphology to have generated debate over their synonmy^8^.

Fabbri et al.^1:Figs. 2, 3^ regress *C*_*g*_ across a broad range of extant and extinct species coded for function (F-flying; D-diving) and frequency (0-unable; 1-able/infrequent; 2-frequent/sustained). For interpreting spinosaurid function, non-flying animals that dive frequently (F = 0, D = 2) are critical for comparison and account for 59 of the taxa considered in the study. Of 21 taxa with highest values of *Cg*, 20 are extinct. Only 16 out of 59 nonflying divers are extant, 4 of which (beaver, tapir, two hippos) are herbivores that do not forage (feed) underwater (see Supplementary information). The scarcity of extant divers with high *Cg* values suggests that density may be enhanced when registered from thin sections of bone in rock. A plot including only extant nonflying divers shrinks parameter space to exclude *Spinosaurus* and *Baryonyx*, in part because of the absence of comparable large-bodied extant divers in the study (Fig. 2).

**Fig. 2.**
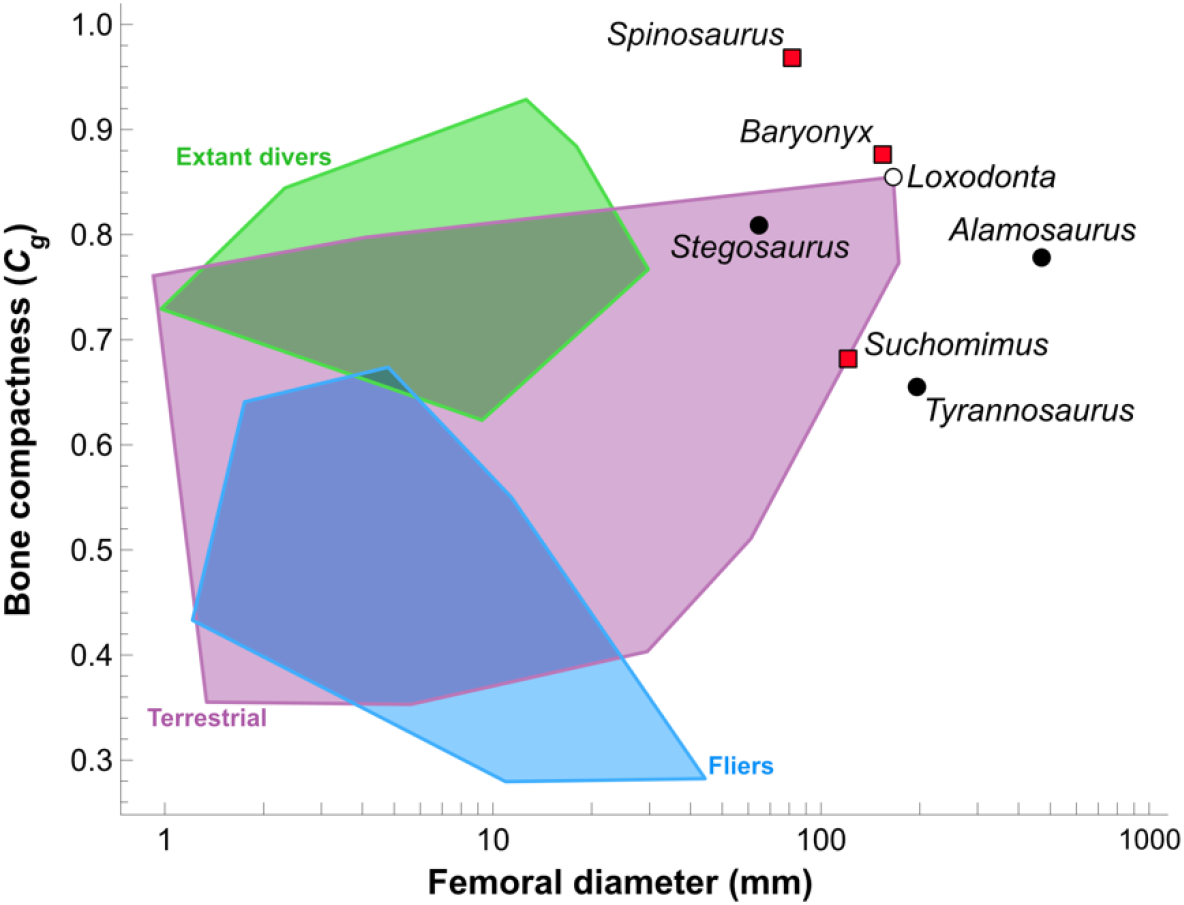
Group overlap and the position of spinosaurids and extant divers. Using the femoral dataset in Fabbri et al.^1^, bone compactness (*Cg*) is regressed on the log of femoral diameter showing the position of the three spinosaurids (*Spinosaurus, Baryonyx, Suchomimus*; red squares) outside the parameter space for “living divers” (n = 16; broad sampling of mammals, crocodylians and birds scored 1 or 2 for diving function; see Supplementary information). The plot shows broad overlap of fliers, divers and terrestrial animals and the relatively compact bone in large extant (elephant, *Loxodonta*; open circle) and extinct (*Stegosaurus, Alamosaurus*; filled circles) quadrupeds and large extinct biped *Tyrannosaurus* (filled circle).

Conversely, Fabbri et al. almost exclusively sample extant taxa for “terrestrial” (i.e., non-flying, non-diving) species (Fig. 2), scoring the diving capacity as “unknown” for 37 extinct nonavian dinosaurs in the dataset (including *Stegosaurus* and desert-living *Oviraptor*). The sample of large ornithischians and sauropods, many of which have infilled limb bones^5,6^, is limited to *Stegosaurus* and one full-sized sauropod (*Alamosaurus*). Both of these, along with the African elephant (*Loxodonta*), have femoral compactness comparable to that of *Baryonyx* (Fig. 2) that would register near the centroid of their violin plot for “nonflying divers.”

### Faulty statistical inferences

Predicting the functional capacity of an extinct species based on bone density requires consistent and unambiguous correlation with function (lifestyle) among extant species, which is not the case in the plots generated by Fabbri et al.^1:Figs. 2, 3^. The terrestrial African elephant (*Loxodonta*) and Madagascar hedgehog (*Tenrec*) plot closest to the regression line for “diving” based on femoral bone density; the terrestrial wombat (*Vombatus*) plots closest to the regression line for “diving” based on rib bone density; and the diving seal (*Phoca*) plots closest to the regression line for “flying” based on rib bone density. Violin plots for femur and rib bone density in *Spinosaurus* and *Suchomimus*, respectively, plot across the distributions of all functional categories.

Broad overlap between groups in the plots of Fabbri et al. precludes their use for inferring the functional categorization of individual extinct species such as spinosaurids (Fig. 2). The problem of *a priori* group membership, in particular, is central to a well-known fallacy in statistical inference (the ecological fallacy)^9^ that occurs when aggregate properties of a group are used to infer the properties of individuals within that group. This includes interpreting an individual species (e.g., *Spinosaurus aegyptiacus*) on the basis of a group distribution, or proximity to a regression line from a group, no matter the quality or quantity of data points. Human male and female body mass, to take well-known case study, broadly overlap and cannot be used to reliably determine the sex of an unknown individual, despite an extraordinarily large, accurate dataset^10^. Fabbri et al. fall prey to this fallacy when they posit behavioral categorizations (diving, terrestrial, flying) for individual extinct species based on generalized linear regressions derived from broadly overlapping distributions.

Body size is a common confounding variable that is not adequately addressed for a dataset of species with body mass spanning several orders of magnitude. Femoral shaft dimensions are strongly correlated with body mass and, as a result, widely used to infer body mass in extinct taxa^11^. Bone density as well as the diameter of medullary space are also correlated with static or dynamic loading^12^. Over-representation of the aquatic reptile *Nothosaurus* in the dataset (6 specimens) shows a range of bone density (0.738-0.938) that is strongly correlated with femoral diameter. The lack of adequate sampling of large terrestrial dinosaurs (ornithischians, sauropods) comparable to *Spinosaurus* leaves open the possibility that its increased bone density is a correlate of large body size.

### Redefinition of the term “aquatic”

“Aquatic,” when applied to a taxon, lifestyle or ecology, is a global characterization with longstanding meaning —an animal that has profound modifications for life in water that markedly constrain life on land (e.g., a dugong)^13,14^. Anything less is “semiaquatic,” no matter the diet, amount of time spent in or near water, or diving capacity. Fabbri et al. discard this longstanding ecomorphological distinction to focus solely on “submergence,” and then use this observed (or inferred) capacity for claiming “aquatic” status. They regard extant hippos as “aquatic” and extant crocodilians as “aquatic archosaurs.” “Fully submerged behavior” (their “subaqueous foraging”) is evidence enough for “aquatic” standing, even when feeding (i.e., foraging) underwater is minimal (crocodylians^15^) or entirely absent (hippos^16^). With such expansive redefinition of this term, humans could pass as “aquatic primates,” given our capacity for diving.

Finally, Fabbri et al.’s claim that previous interpretations of the function, or lifestyle, of nonavian dinosaurs largely excluded shoreline, shallow water habitats is a straw man. Ever since its discovery in 1915, *Spinosaurus* has been viewed as a specialized shoreline piscivore^17^, and ensuing years have witnessed accumulating footprint evidence that documents wading or swimming dinosaurs from the major clades^18^. Discerning the functional capacities or specific lifestyle of these species in water will require better controlled comparative histology, quantitative whole-body comparisons to aquaphilic vertebrates, and especially biomechanical modeling^19^.

## Data Availability

All data are available in this contribution and its Supplementary Information.

## Author contributions

N.M. and P.C.S. wrote the initial draft, S.L.B., D.V., N.M. and P.C.S collaborated on the figures, and K.K.F., D.V., F.E.F. and D.M.H. made substantive edits and contributions to the final version.

## Additional information

### Supplementary information

The online version contains supplementary available at https://*****

