## Supplemental Information for "Spinosaurids as ‘subaqueous foragers’ undermined by selective sampling and problematic statistical inference"

ARISING FROM **M. Fabbri et al. nature 9https://doi.org/10.1038/s41586-022-04528-0**

[1]Intellectual Ventures, 3150 139th Avenue Southeast, Bellevue, WA 98005.
[2]Department of Organismal Biology, University of Chicago, Chicago, IL, 60637.
[3]Committee on Evolutionary Biology, University of Chicago, Chicago, IL, 60637.
[4]Department of Earth Sciences, University of Southern California, Los Angeles, CA 90089.
[5]Dinosaur Institute, Natural History Museum of Los Angeles County, Los Angeles, CA 90007.
[6]Grupo de Biología Evolutiva, UNED, 28040 Madrid, Spain.
[7]Department of Biology, West Chester University, West Chester, PA 19383.
[8]Royal Tyrrell Museum of Palaeontology, Drumheller, Alberta, Canada, TOJ OY.

**Contents**

1. Bone density metrics
2. Femoral form and thin section in *Spinosaurus aegyptiacus* and *Suchomimus tenerensis*
3. Sampling/categorization issues
4. Confounding factors for Cg signal
5. Statistical fallacies in ascertaining lifestyle
   Figs. S1-5, Tables S1, S2
6. Materials
7. Acknowledgments

### 1. Bone density metrics and diameter

*Bone Profiler* estimates bone compactness (the relative area of a cross-section occupied by mineralized bone tissue) by calculating the proportion of bone area observed in 51 zones averaged across 60 radial sectors, and it constructs a "compactness profile" from the centroid to the periphery of the outer bone wall[1,2]. Several parameters of the compactness profile can be used to describe the distribution of bone mass within a cross-section.

The metric used by Fabbri et al.[3] as a proxy for bone density, global bone compactness (*Cg*), is the ratio between surface covered by mineralized tissues and section area, one of many metrics for compactness. Over the last decade, other metrics generated by this program (*S*, *P*, *Min, Maxrad, Cc, Cp*) were shown to be better correlated with lifestyle based on extant amniotes[4,5]. Because the plots in Fabbri et al.[3:Figs. 2,3] include several examples where extant taxa are projected to have lifestyles strongly at variance with their documented lifestyle, the dataset should be replotted using a more effective suite of bone density metrics.

Fabbri et al. do not provide details of how they assembled the set of bone compactness data from prior studies, or how they calibrate any potential differences between data measured from CT scans versus thin sections.

The data also include "diameter" but there is no description of how this is defined for bones that are not circular in cross section.

### 2. Femoral form and thin section in *Spinosaurus aegyptiacus* and *Suchomimus tenerensis*

The femur in *Spinosaurus aegyptiacus* differs from that in baryonychine spinosaurids in several ways (Fig. S1). The femur is shorter that the tibia, a proportion otherwise associated with speed in theropods, and the fourth trochanter is hypertrophied, extending distally to mid shaft (Fig. 2c). As a result, only a short portion of the femoral shaft in the distal half of the bone has a subcircular cross section. One of us (PCS) made a histologic cross section in that portion of the shaft[6] (used in Fabbri et al.[3]), which shows the absence of a substantial medullary cavity (Fig. S1b).

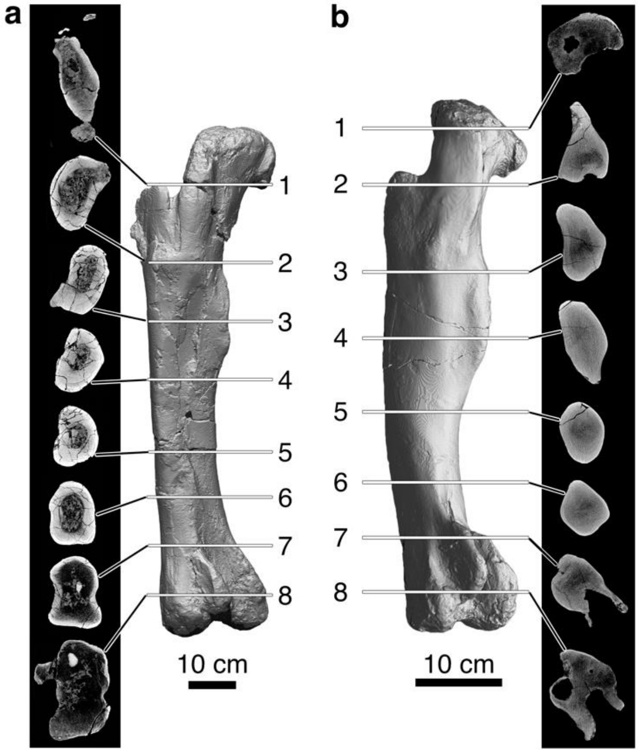


**Fig. S1 Spinosaurid femoral form and cross sectional density a**, Left femur of *Suchomimus tenerensis* (MNBH GAD500, holotype, length 107.5 cm) in posterolateral view with eight spaced CT cross sections. **b**, Left femur of *Spinosaurus aegyptiacus* (FSAC-KK 11888, neotype, length 61.0 cm) in posterolateral view with eight spaced CT cross sections. FSAC, Faculté des Sciences Aïn Chock, University of Casablanca, Morocco; MNBH, Musée National Boubou Hama, Niamey.

A second femur of *Spinosaurus* cf. *aegyptiacus*[7] (Figs. 1a,b, S2) preserves approximately 47% of the proximal end of the femur, judging from the complete femora of the neotypic skeleton. The second femur has an estimated complete length of ~67 cm, which is very close to the preserved length (61.0 cm) of the most complete neotypic femur (Fig. S1b). The proximal shaft of the second femur, unlike the neotypic femur, has a narrow medullary cavity lined by cancellous bone that would have extended at least to its midshaft. A CT scan of the neotypic femur shows less dense bone in the same general region of the medullary cavity in the second femur (Fig. S1b). The presence of a medullary cavity would have lowered global bone compactness.

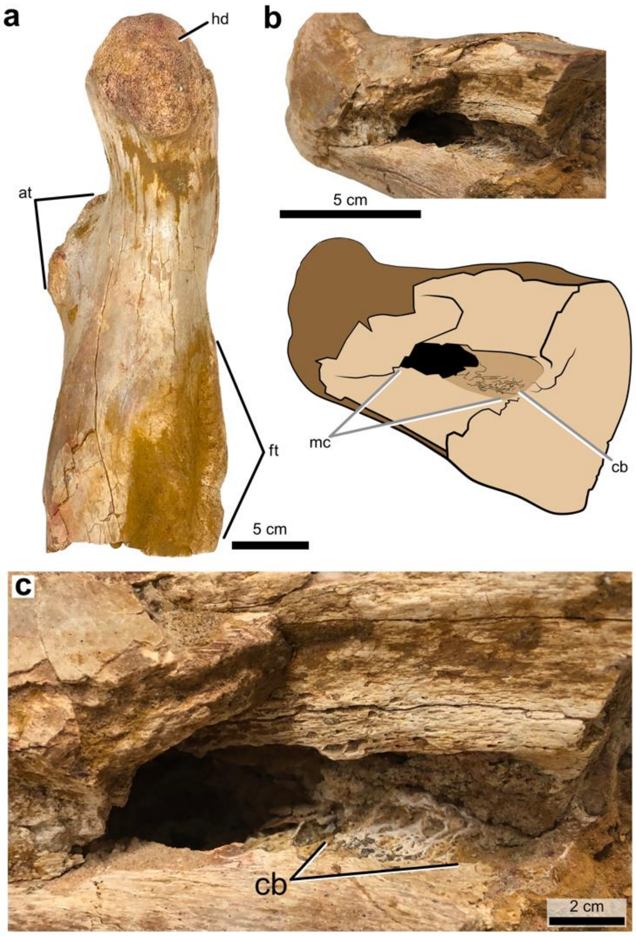


**Fig. S2** ***Spinosaurus cf. aegyptiacus* femur. a**, Proximal half of the right femur (NMC 41869) in medial view. **b**, Medullary cavity in ventrolateral view. **c**, Bone lining the medullary cavity. National Museum of Canada, Ottawa. at, anterior trochanter; cb, cancellous bone; ft, fourth trochanter; hd, head; mc, medullary cavity.

Fabbri e al.[1:Suppl. Fig. S2] figure two magnified thin sections identified as *Suchomimus tenerensis* identified as “G51” and “G94,” which are field numbers for the holotype (MNBH GAD500) a referred subadult individual (MNBH GAD70), respectively. Neither of these specimens have been sectioned, however, and we are unable to match those thin sections to the thin section made by one of us (PCS) of an adult femur of *Suchomimus tenerensis* (MNBH GAD99), which was also figured by Fabbri et al. (Fig. S3d). Their figure, nonetheless, differs from a photograph of the thin section (Fig. S3b), which shows significantly more cancellous bone toward the center of its shaft and a smaller medullary cavity (Fig. S3b, c). This section was taken from the distal end of the shaft of an adult individual (Fig. S3a) of the same size as the holotype (both have distal condyles measuring 23 cm in width). We show here an additional section for *Suchomimus tenerensis* pertaining to a juvenile individual with femur length approximately half of adult length (Fig. S3e). The medullary cavity of the juvenile is relatively large and surely decreases in relative diameter with growth.

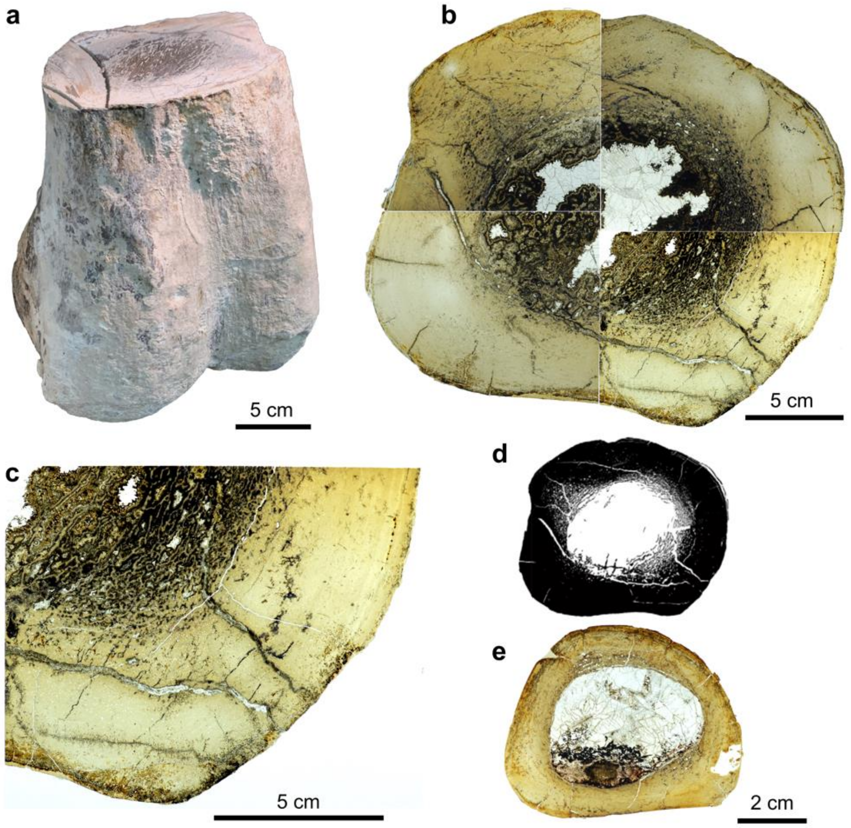


**Fig. S3 *Suchomimus tenerensis* adult and juvenile femoral thin sections. a**, Distal end of an adult right femur with an estimated length of ~1.0 m. (MNBH GAD99). **b**, Thin section of the distal shaft (MNBH GAD99). **c**, Magnified view of the thin section showing cancellous bone surrounding the medullary cavity (MNBH GAD99). **d**, Depiction of the thin section in Fabbri et al.[3:Fig. 1b] **e**, Thin section of the midshaft of a right femur of a juvenile individual (femur length 55.3 cm; MNBH GAD72). MNBH, Musée National Boubou Hama, Niamey.

**3. Sampling/categorization issues**

Fabbri et al. argue that bone compactness (Cg) is a signal of secondary aquatic lifestyle in both extinct and extant taxa and across a wide range of amniotes. Subsets of the dataset compiled by Fabbri et al.[3] contain a number of anomalies, however, that appear to have skewed their results. These data subsets involve:

(1) Femoral bone density in 59 taxa representing "nonflying divers" (abbreviated F=0, D=2) (Table S1);

(2) Femoral bone density in taxa interpreted as terrestrial (F= 0, D=0); and

(3) Femoral bone density in 37 extinct nonavian dinosaurs (F= 0, D=unknown) (Table S2).

The composition of these three subsets is strongly affected by sampling bias (extinct vs extant) and the scoring of function/lifestyle categorization (positively scored vs unknown). Both compromise comparisons and favor their conclusions regarding "subaqueous foraging."

**Extant vs extinct**

The "nonflying diver" dataset, for example, is composed overwhelmingly of extinct taxa (43, 71.7%). The smaller number of extant taxa in this subset (16, 26.7%) include four herbivores (e.g., hippo) that feed on terrestrial plants and, thus, cannot be regarded as "subaqueous foragers"

(Table S1). This important subset of data ("nonflying divers") is better limited to 43 extinct (78.2%) and 12 extant (21.8%) taxa.

These 43 extinct taxa are strongly biased toward the highest values of Cg. Nineteen (95%) of the top 20 taxa ranked by Cg (Cg > 0.89) are extinct. At face value, this suggests that high bone compactness is somehow time limited and generally lower among extant species. Or this may represent systematic error in how compactness is measured optically from fossilized bone infilled with rock matrix resulting in increased Cg. Or this is simply skewed sampling. A simple Monte Carlo simulation shows that it would be highly unlikely (< 1% probability) that 43 extinct and 12 extant taxa, respectively, would be randomly drawn from equivalent distributions of Cg would yield Cg values this skewed. Fabbri et al. use PGLS to correct for phylogenetic proximity among sampled taxa, although this method does not compensate for skewed sampling that favors one subset of taxa over another. In particular it cannot compensate for taxa that were omitted from the analysis.

The "terrestrial" femoral dataset (F=0, D=0), in stark contrast to "nonflying divers," comprises 59 taxa that are overwhelmingly extant (54 taxa, 91.5%), with few extinct taxa (5, 8.5%). High values of Cg are necessarily dominated by extant taxa, which have lower Cg (< 0.855) than that of *Spinosaurus* (0.968). No reason is given for the stark difference in sampling between these first two cohorts of taxa, the net effect of which is to lower Cg for terrestrial taxa compared to aquatic foragers.

**Known vs unknown**

Function/lifestyle categorization is also inexplicably skewed in terms of what is considered 'knowable' among extinct taxa. "Nonflying divers" include many extinct species, all of which are scored as "subaqueous foragers" (F=0, D=2). The 37 nonavian dinosaurs in the "terrestrial" subset, in contrast, are scored as nonflying reptiles of "unknown" capacity regarding diving (F=0, D=unknown) (Table S2). All included nonavian dinosaurs, in other words, potentially could be "divers," including those with elephantine feet and extensive armor (e.g., *Stegosaurus*), those living in xeric habitats (e.g., *Oviraptor*), and those of enormous body size unearthed from inland terrestrial deposits (e.g., *Alamosaurus*). Fabbri et al. are able to discern and score diving function for one set of extinct taxa ("aqueous foragers") but find it impossible to evaluate and score diving function for the extinct ("terrestrial") nonavian dinosaurs. This seems arbitrary. Most of the nonavian taxa in the study have always been considered fully terrestrial, based on a suite of palaeontologic and geologic evidence. Although some sauropods were once considered semiaquatic, their fully terrestrial lifestyle has been a settled matter since the mid 1980s[8].

Many "nonflying divers" in the study, such as ichthyosaurs, mosasaurs and cetaceans, have reduced or vestigial limbs, profound adaptations that render an aquatic lifestyle an obvious interpretation. However, this begs the question: why should these highly derived, aquatically adapted taxa be included as suitable comparative models for terrestrial limbed spinosaurids? That forces a false choice that offers for comparison with spinosaurids either fully aquatic extinct taxa (with high Cg) or terrestrial extant taxa (with lower Cg). This alone seems sufficient to force the outcome of the analysis.

**Femoral vs rib density signal**

For Cg to stand as a reliable measure of secondary aquatic lifestyle related to buoyancy as the authors claim, femoral and rib datasets in Fabbri et al. should give broadly congruent results. They do not. The rib data for "nonflying divers" (F=0, D=2) comprises 49 taxa evenly split between extinct (25 taxa, 51%) and extant (24 taxa, 49%) species. Ranking them by Cg shows no bias favoring either extant or extinct taxa, and their average Cg is not markedly higher that other categories of function/lifestyle. Why would the density in one bone (femur) register function/lifestyle and not another (rib)? Bone density and its effect on buoyancy among living and extinct amniotes is fraught with complexity, variation and evolutionary reversal, requiring very careful comparative analysis to prove its adaptive presence in all but the extreme cases involving pachystosis[9].

**Table S1. Nonflying, diving subsample of taxa.** The taxa used in the analysis of bone density by Fabbri et al.[3] include 59 that are scored as "nonflying divers" (flying F = 0, diving D = 2), which we rank below by their Cg value. Extant species are indicated (grey shade) and marked as to whether they forage (feed) underwater. The six specimens of *Nothosaurus* are in bold.

| No. | Taxon | Flying | Diving | Femoral diameter | $C_g$ | Extant (E) | Extant Subaqueous Forager |
|---|---|---|---|---|---|---|---|
| 1 | *Serpianosaurus* | 0 | 2 | 4.8 | 0.989 | — | — |
| 2 | large_Eocene_stem_penguin | 0 | 2 | 16.744 | 0.988 | — | — |
| 3 | *Maiacetus* | 0 | 2 | 30.43 | 0.985 | — | — |
| 4 | *Nanophoca_vitulinoides* | 0 | 2 | 20.3 | 0.973 | — | — |
| 5 | *Cryptoclidus* | 0 | 2 | 84.08 | 0.97 | — | — |
| 6 | *Champsosaurus_* | 0 | 2 | 7.85 | 0.968 | — | — |
| 7 | *Neusticosaurus* | 0 | 2 | 19.1 | 0.968 | — | — |
| 8 | *Phocanella_pumila* | 0 | 2 | 29.5 | 0.966 | — | — |
| 9 | *Placondontia_*indet | 0 | 2 | 23.38 | 0.959 | — | — |
| 10 | ***Nothosaurus*_102** | 0 | 2 | 5.168 | 0.955 | — | — |
| 11 | *Champsosaurus* | 0 | 2 | 12.389 | 0.952 | — | — |
| 12 | Small_Eocene_penguin | 0 | 2 | 9.457 | 0.942 | — | — |
| 13 | *Paraplacodus* | 0 | 2 | 9.05 | 0.939 | — | — |
| 14 | ***Nothosaurus*_150** | 0 | 2 | 8.125 | 0.938 | — | — |
| 15 | *Rhaeticosaurus* | 0 | 2 | 36 | 0.936 | — | — |
| 16 | *Caiman_yacare* | 0 | 2 | 12.623 | 0.929 | E | yes |
| 17 | *Basilosaurus* | 0 | 2 | 21.96 | 0.926 | — | — |
| 18 | ***Nothosaurus*_568** | 0 | 2 | 5.464 | 0.909 | — | — |
| 19 | *Anarosaurus* | 0 | 2 | 10 | 0.901 | — | — |
| 20 | *Plesiosaurus* | 0 | 2 | 41 | 0.9 | — | — |
| 21 | *Rodhocetus* | 0 | 2 | 26.863 | 0.893 | — | — |
| 22 | *Desmana_moschata* | 0 | 2 | 5.1 | 0.89 | E | yes |
| 23 | *Alligator* | 0 | 2 | 18 | 0.884 | E | yes |
| 24 | *Cricosaurus* | 0 | 2 | 16.265 | 0.874 | — | — |
| 25 | *Spheniscus_humboldti* | 0 | 2 | 8.06 | 0.872 | E | yes |
| 26 | *Ornithorhynchus_anatinus* | 0 | 2 | 5.21 | 0.871 | E | yes |
| 27 | *Indohyus* | 0 | 2 | 7.44 | 0.867 | — | — |
| 28 | *Simosaurus* | 0 | 2 | 22.97 | 0.865 | — | |
| 29 | *Aptenodytes* | 0 | 2 | 16.395 | 0.864 | E | yes |
| 30 | *Placodontia_*indet_1 | 0 | 2 | 20.97 | 0.859 | — | — |

| | | | | | | | |
|---|---|---|---|---|---|---|---|
| 31 | *Lutra_vulgaris* | 0 | 2 | 10.02 | 0.85 | E | yes |
| 32 | *Chironectes_minimus* | 0 | 2 | 4.78 | 0.849 | — | — |
| 33 | *Pistosaurus* | 0 | 2 | 27.56 | 0.845 | — | — |
| 34 | *Micropotamogale_euwenzorii* | 0 | 2 | 2.31 | 0.844 | E | yes |
| 35 | *Psephoderma* | 0 | 2 | 9.37 | 0.843 | — | — |
| 36 | *Metryorhynchus* | 0 | 2 | 27.384 | 0.828 | — | — |
| 37 | ***Nothosaurus_mirabilis*** | 0 | 2 | 16.09 | 0.828 | — | — |
| 38 | *Hippopotamus_amphibius* | 0 | 2 | 59.34 | 0.828 | E | no[14] |
| 39 | *Otaria_byronia* | 0 | 2 | 22.28 | 0.821 | E | yes |
| 40 | *Palaeospheniscus* | 0 | 2 | 8.52 | 0.792 | — | — |
| 41 | *Ichtyosaur_sp.* | 0 | 2 | 165.44 | 0.776 | — | — |
| 42 | ***Nothosaurus_mirabilis_1*** | 0 | 2 | 21.7 | 0.776 | — | — |
| 43 | *Choeropsis_liberiensis* | 0 | 2 | 29.78 | 0.767 | E | no |
| 44 | *Remingtonocetus* | 0 | 2 | 35.72 | 0.765 | — | — |
| 45 | *Simosaurus_1* | 0 | 2 | 22.9 | 0.764 | — | — |
| 46 | *Castor_fiber* | 0 | 2 | 29 | 0.749 | E | no[15] |
| 47 | ***Nothosaurus_giganteus*** | 0 | 2 | 26.819 | 0.738 | — | — |
| 48 | *Callophoca_obscura* | 0 | 2 | 25.86 | 0.733 | — | — |
| 49 | *Leptophoca_proxima* | 0 | 2 | 28.9 | 0.729 | — | — |
| 50 | *Neomys_fodiens* | 0 | 2 | 0.969 | 0.729 | E | yes |
| 51 | *Hexaprotodon_garyam* | 0 | 2 | 69.4 | 0.726 | — | — |
| 52 | *Hesperornis* | 0 | 2 | 22.914 | 0.725 | — | — |
| 53 | *Hydromys_chrysogaster* | 0 | 2 | 5.42 | 0.689 | E | yes |
| 54 | *Tapirus_terrestris* | 0 | 2 | 33.2 | 0.687 | E | no[16] |
| 55 | *Protochampsidae* | 0 | 2 | 10.17 | 0.673 | — | — |
| 56 | *Ichtyosaurus_* | 0 | 2 | 86.48 | 0.659 | — | — |
| 57 | *Dyrosaurid* | 0 | 2 | 12.54 | 0.635 | — | — |
| 58 | *Phalacrocorax_harrisi* | 0 | 2 | 9.26 | 0.623 | E | yes |
| 59 | *Desmostylus_hesperus* | 0 | 2 | 38 | 0.596 | — | — |

**Table S2. Femoral Cg data from nonflying taxa with unknown diving ability (F=0, D= unknown).** Fabbri et al.[3] included a subset of 37 extinct nonavian dinosaurs, which we rank by their Cg value. These taxa are classified as "unknown" with respect to diving ability, despite exhibiting skeletal anatomy consistent with a terrestrial lifestyle. There are three spinosaurids (red) and two sauropods (bold).

| No. | Taxon | Flying Score | Diving Score | Femur Diameter | Cg |
|---|---|---|---|---|---|
| 1 | *Spinosaurus* | 0 | Unknown | 81.52 | 0.968 |
| 2 | *Baryonyx* | 0 | Unknown | 154 | 0.876 |
| 3 | *Stegosaurus_sp* | 0 | Unknown | 64.3 | 0.81 |
| 4 | ***Alamosaurus*** | 0 | Unknown | 470.2 | 0.777 |
| 5 | *Dysalotosaurus* | 0 | Unknown | 36.297 | 0.733 |

| | | | | | |
|---|---|---|---|---|---|
| 6 | *Torvosaurus* | 0 | Unknown | 132.57 | 0.728 |
| 7 | *Tenontosaurus* | 0 | Unknown | 22.02 | 0.714 |
| 8 | Unamed_theropod_kem_kem | 0 | Unknown | 22.37 | 0.705 |
| 9 | ***Antetonitrus*** | 0 | Unknown | 43.97 | 0.702 |
| 10 | *Gorgosaurus* | 0 | Unknown | 46.9 | 0.689 |
| 11 | *Tyrannotitan* | 0 | Unknown | 156.83 | 0.682 |
| 12 | *Suchomimus* | 0 | Unknown | 120.6 | 0.682 |
| 13 | *Megalosaurus* | 0 | Unknown | 118.21 | 0.678 |
| 14 | *Scutellosaurus_lawleri* | 0 | Unknown | 12.5 | 0.671 |
| 15 | *Mussaurus* | 0 | Unknown | 110.42 | 0.67 |
| 16 | *Troodon_formosus* | 0 | Unknown | 27.822 | 0.665 |
| 17 | *Tyrannosaurus* | 0 | Unknown | 197 | 0.656 |
| 18 | *Deinonychus* | 0 | Unknown | 37.972 | 0.632 |
| 19 | *Gallimimus*_sp | 0 | Unknown | 36.9 | 0.626 |
| 2 | *Fruitadens* | 0 | Unknown | 4.4 | 0.603 |
| 21 | *Allosaurus* | 0 | Unknown | 89.498 | 0.597 |
| 22 | *Plateosaurus* | 0 | Unknown | 67.1 | 0.591 |
| 23 | *Syntarsus* | 0 | Unknown | 25.3 | 0.58 |
| 24 | *Rativates* | 0 | Unknown | 33.35 | 0.572 |
| 25 | *Masiakasaurus* | 0 | Unknown | 19.95 | 0.567 |
| 26 | *Condorraptor* | 0 | Unknown | 63.312 | 0.567 |
| 27 | *Asilisaurus* | 0 | Unknown | 8.414 | 0.565 |
| 28 | *Gobiraptor_* | 0 | Unknown | 27.3 | 0.563 |
| 29 | *Australovenator* | 0 | Unknown | 77.186 | 0.56 |
| 30 | *Lepidus* | 0 | Unknown | 27.42 | 0.551 |
| 31 | *Oviraptor_caenagnathid* | 0 | Unknown | 32.25 | 0.499 |
| 32 | *Halszkararaptor_* | 0 | Unknown | 7.617 | 0.493 |
| 33 | *Asfaltovenator* | 0 | Unknown | 103.03 | 0.47 |
| 34 | *Eustreptospondylus* | 0 | Unknown | 50.1 | 0.459 |
| 35 | *Avimaia* | 0 | Unknown | 1.75 | 0.458 |
| 36 | *Oviraptor* | 0 | Unknown | 27.769 | 0.438 |
| 37 | noasaurid_kem_kem | 0 | Unknown | 22.25 | 0.393 |

### 4. Confounding factors for Cg signal

Fabbri et al. argue that the bone compactness metric Cg is a reliable signal for secondary aquatic lifestyle involving prolonged submergence across amniotes. To the extent that Cg is correlated with other unrelated factors, correlation with aquatic lifestyle is compromised. The proportion of extinct versus extant taxa discussed above is potentially confounding for Cg if these categories show consistent differences.

Burrowing is a function known to increase Cg[10], and several fossorial taxa are present in the dataset. Aquatic lifestyle increases Cg as well, although some taxa have secondarily reduced Cg as an aquatic adaptation[9]. Another possible confounding factor is body size, which can be demonstrated by plotting Cg in the overrepresented genus *Nothosaurus*. The dataset includes six specimens of *Nothosaurus*, and three specimens of two closely related taxathat show a strong inverse correlation between Cg and body size (Fig. S4). the sources for these data span four different studies in the literature, so it is not caused by a problem in a single study.

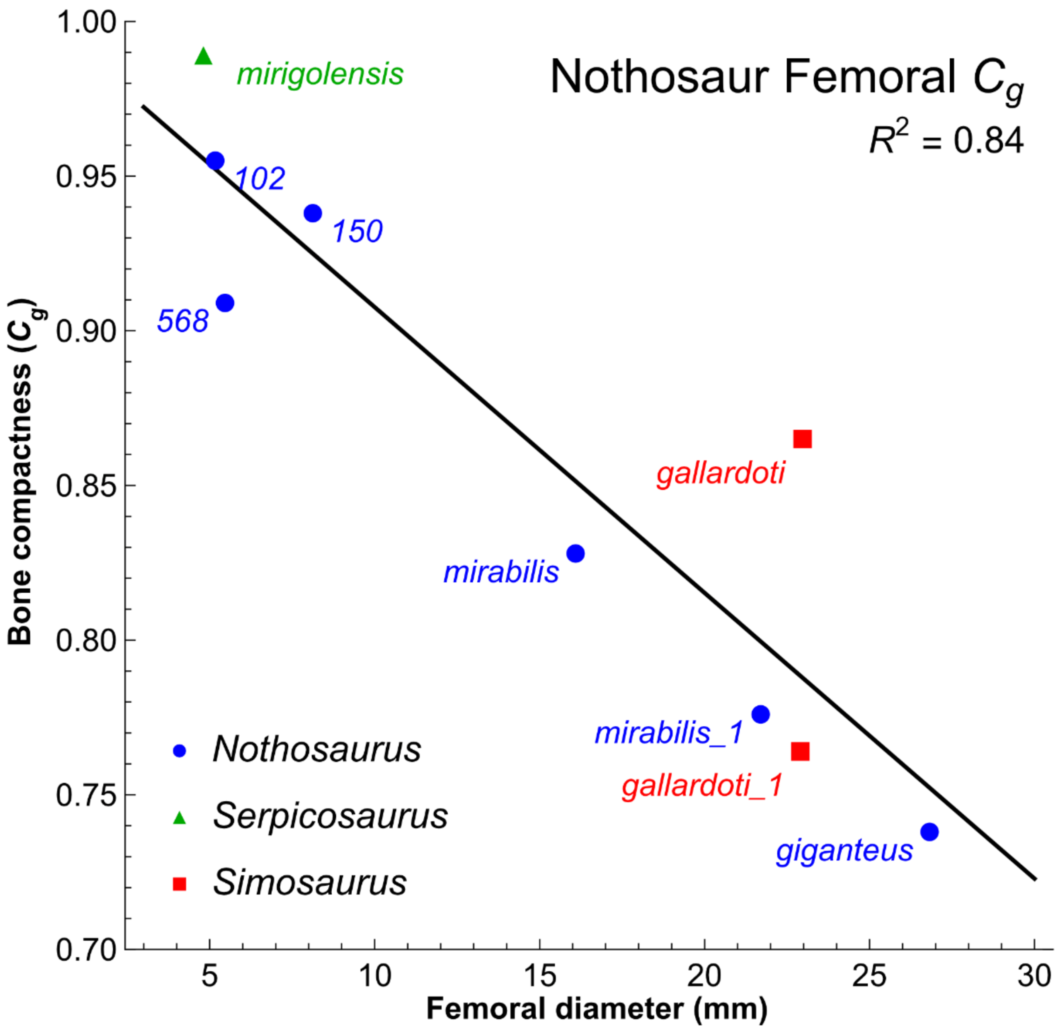


**Fig. S4 Correlation between global bone compactness and body size in Nothosaurid and Pachypleurosaurid marine reptiles.** Fabbri et al.[3] include six specimens of *Nothosaurus* , two of the related northosaur *Simosaurus* and one of the related pachypleurosaur *Serpicosaurus*. They show a very strong inverse correlation between global bone compactness (Cg) and femoral diameter, a common proxy for body size. Larger diameter femora correlate with lower Cg (for whatever reason), strongly suggesting that body size exists as a potential confounding factor in sampled taxa.

**5. Statistical fallacies in ascertaining lifestyle**

The example we cited using adult human body weight[11] illustrates the dangers of the ecological fallacy[12]. Although the average male is heavier than the average female, overlap in the distributions eliminates the possibility of using the group-wide average or other properties derived from the aggregate (e.g., equiprobable mass) for identifying the proper classification of an outside individual (Fig. S5). This approach to classification would only work with high accuracy if the distributions had little overlap.

To determine the accuracy of classification, we must fit a smooth distribution to the histogram data (extreme value distribution) and find the body mass which is equally probable in each distribution. The histograms are well modeled by the extreme value distribution (center panel), and the equiprobable point occurs as a mass of 72.46 kg (right panel). This probability-based method will classify an individual specimen of unknown group membership into the groups male or female depending on whether they are above or below the equiprobable point of 72.46 kg. Overlap between the distributions causes errors about 38% of the time – either for males below 72.46 kg, or females above that value, assuming there is an equal probability of the unknown subject being male or female A random guess of the group will be in error 50% of the time, so using body mass to classify in this manner is only a ~12% improvement.

Previous studies examining the relationship between bone compactness and lifestyle use linear discriminant analysis[2], a clustering technique that attempts to find a line or plane in parameter space separating clusters of data points. This method and other suitable alternatives, such as logistic regression, do not work when there broad overlap between data partitions[13].

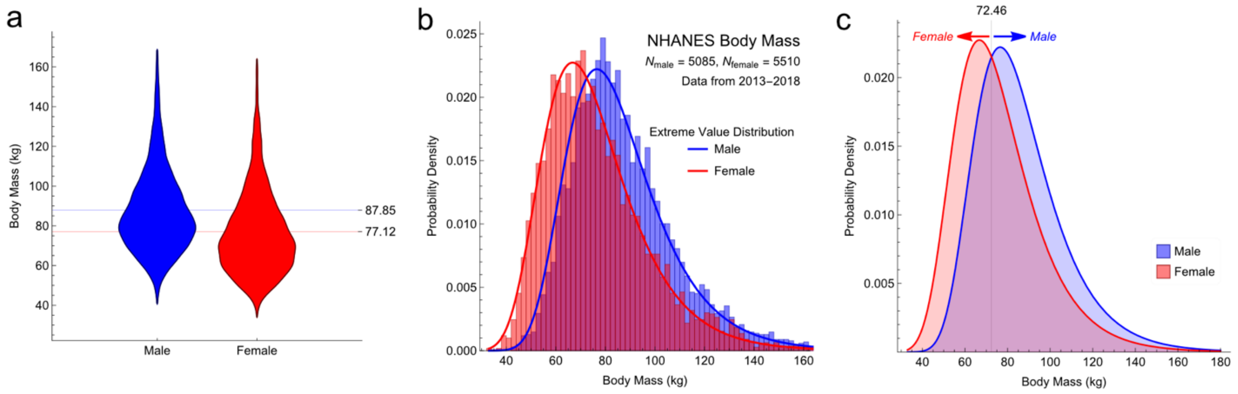


**Figure S5. The 'ecological fallacy' using data on human adult body mass. a,** Violin plot of body mass distributions. **b**, Histogram of body mass distributions with fits to the extreme value distribution. **c**, Probability curve for classification of sex by body mass.

Failure to classify an individual by comparison comes from using group-wide aggregate measures such as the distribution. Statistics provide many analytical tools (mean, variance, and other tools) for comparing a sample group to another group to assess whether they were drawn from the same underlying population, such as Student's t-test, Welch's t-test, the Mann-Whitney test and the F and Z tests. When sample group size (N) is small, these methods provide low statistical power. Classifying a single human, or in Fabbri et al. *Spinosaurus* is an N = 1 example, wherein these statistical tools simply do not work. The ad hoc method of Fabbri et al. has no support in the statistical literature.

In large human study alluded to above[11], group composition was carefully chosen to accurately model population-wide adult human body mass. Data points potentially contaminated with confounding contributions to body mass were eliminated (e.g., removing pregnant women). The groups identified by Fabbri et al.[3], in contrast, have not been controlled for confounding factors such as body mass, burrowing and other factors known to contribute to bone compactness, and only two categorical variables (flying, diving) are considered, which forces an overly constrained choice of only two possibilities ("subaqueous forager" versus not).

The analysis of Fabbri et al.[3] is more complex that the human example, because they use PGLS-based linear regression models rather than simple distributions to process bone density data in order to reduce potentially confounding effects of phylogenetic signal. This does not avoid the ecological fallacy, because the comparison group is still N=1 for *Spinosaurus*. If one includes all spinosaurids for a group of N = 3, statistical power remains weak. This larger grouping also assumes incorrectly that all spinosaurids have equal degrees of secondary aquatic function.

### 6. Materials

The CT scans for the FSCA-KK 11888 and MNBH GAD500 femora used in this reply are available for review on Morphosource at the following URLs:

*Spinosaurus* femur FSCA-KK 11888:
https://www.morphosource.org/concern/media/000434660?locale=en
*Suchomimus* femur MNBH GAD500:
https://www.morphosource.org/concern/media/000434666?locale=en

### 7. Acknowledgments

We thank the fossil preparators of the Sereno lab for fine preparation of the FSAC-KK 11888 femur and for preparation of the MNBH GAD500 femur. We also thank Nicholas Gruszaukas, David Klein, and the University of Chicago Department of Radiology for scanning those femora. Wayt Gibbs provided editorial assistance.

12. Piantadosi, S., Byar, D. P. & Green, S. B. The ecological fallacy. *Am. J. Epidem.* **127**, 893–904 (1988).
13. Hastie, T., Tibshirani, R. & Friedman, J. *The Elements of Statistical Learning. Data Mining, Inference, and Prediction* (second ed.). Springer. 745 pp. (2009).
14. Owen-Smith, R. N. *Megaherbivores: The Influence of Very Large Body Size on Ecology*. (Cambridge Univ. Press, 2000).
15. Haarberg, O. & Rosell, F. Selective foraging on woody plant species by the Eurasian beaver (*Castor fiber*) in Telemark, Norway. *J. Zool.* **270**, 201–208 (2006).
16. Fruit patch size and frugivory in the lowland tapir (*Tapirus temstris*). *J. Zool.* **222**, 121–128 (1990).
